# Historically Small Population Size Limits Purging of Deleterious Mutations in a Conservation-Reliant Species, the Kirtland’s Warbler

**DOI:** 10.64898/2026.05.15.725193

**Authors:** Anna María Calderón, Alexander T. Salis, David P.L. Toews, Zachary A. Szpiech

## Abstract

Strong population contractions can leave a persistent genomic legacy that can influence populations long after their demographic recovery. While bottlenecks facilitate the removal of strongly deleterious mutations, the effectiveness of purging may be limited in historically small populations. The Kirtland’s warbler (*Setophaga kirtlandii*) is a rare North American songbird with an ancestrally small population. After narrowly evading extinction, they are one of few species that have been delisted from federal protections in the USA. Despite their recovery, a previous study showed evidence for recent inbreeding and a high burden of deleterious mutations that may have not been purged despite strong bottlenecks. Historical DNA offers a unique opportunity to understand the consequences of recent demographic declines on genetic diversity. Here, we use DNA from over 100-year-old museum specimens to estimate changes in genetic load in the Kirtland’s warblers pre- and post-bottleneck. We validate our results with forward-in-time genetic simulations and explore how sample size and missing data can affect estimates. Both empirical data and simulations suggest a reduced ability to purge deleterious mutations in this historically small population. Our simulations also highlight that limited sampling design and data quality can constrain the ability to detect changes.

## INTRODUCTION

Anthropogenic activities have led to alarming declines in wildlife populations worldwide, with birds and mammals being among the most substantially impacted groups (Ceballos et al., 2015; Rosenberg et al., 2019). Increasing threats from habitat loss, climate change, pollution, and exploitation are becoming more urgent (Capdevila et al., 2026), especially amid weakening environmental protections and insufficient climate policy. In addition to declines, populations are also burdened with increased genomic erosion as it lowers mean population fitness and reduces adaptive potential to future environmental change (Femerling et al., 2023; Mathur et al., 2023). Population size is a key driver of evolutionary processes, influencing the strength of selection, shaping genetic diversity, and the composition of genetic load.

While changes in genetic load can impact the mean fitness of individuals and populations, its relationship to extinction risk remains unclear and contentious (Kardos et al., 2024; Kyriazis et al., 2021). As a metric, genetic load summarizes the frequencies of deleterious alleles across loci, weighted by their selection and dominance coefficients and it is often described in terms of its individual components: the masked and realized load (Bertorelle et al., 2022). The masked load refers to the portion of genetic load that is hidden from selection in heterozygotes but can become realized via inbreeding as these alleles become homozygous. While population bottlenecks can change the constitution of genetic load, the realized portion may also be eliminated depending on the efficacy of selection operating in the population (Dussex et al., 2023). The purging of deleterious mutations, however, may come at a cost via inbreeding depression and the severity can depend on the long-term evolutionary history of the population. For example, ancestrally large populations tend to harbor a greater masked load that poses a greater risk to individual fitness when deleterious alleles are exposed through population bottlenecks and subsequent inbreeding (Femerling et al., 2023; Hedrick & Garcia-Dorado, 2016; Kyriazis et al., 2021). In contrast, ancestrally small populations are thought to be buffered against severe inbreeding depression as their masked genetic load has been largely purged slowly over time (Mathur et al., 2023). Despite this “genetic insurance”, historically small populations can still experience inbreeding depression driven by the cumulative effects of many weak to moderately deleterious mutations (Kardos et al., 2021; Stoffel et al., 2021).

The effects of population history on genetic load dynamics have been a subject of interest and debate for many years (Lande, 1994; Lynch & Gabriel, 1990; Robinson et al., 2018; Simons et al., 2014; Simons & Sella, 2016; van der Valk et al., 2019). Understanding of these dynamics requires the identification of the deleterious mutations that make up genetic load. In the genomics era, numerous studies have used bioinformatic approaches to identify deleterious mutations from sequencing data. However, these methods are inherently limited in their ability to classify such mutations, as they do not provide information on their selection or dominance coefficients (Kardos et al., 2024). Consequently, direct measures of genetic load are currently unavailable for wild populations. In pursuit of this, the R_A/B_ statistic (Xue et al., 2015) is commonly used in conservation genetics to track changes in the genetic load in one population relative to another (Dussex et al., 2021; Grossen et al., 2020; Robinson et al., 2018). To this end, incorporating historical data into conservation genomic analyses can help reveal how recent anthropogenically driven population declines have directly influenced genetic load. Here, we take advantage of museum samples in 100-year-old specimens to explore genetic load dynamics in a conservation-reliant species—a species with an ancestrally small population that experienced a prolonged bottleneck followed by recent recovery.

Kirtland’s warblers (*Setophaga kirtlandii*) are a rare North American songbird with a small breeding distribution largely constrained to the Lower Northern Peninsula of Michigan (**Figure 1A**). This limited distribution is ecologically driven by an obligatory relationship with early successional Jackpine forests (Probst, 1986). Changes in land management, namely the suppression of large-scale wildfires and agricultural expansion, led to a major loss of breeding habitat in this species. After narrowly evading extinction in the 1970s, populations recovered following decades of intense conservation management (Fish and Wildlife Service, 2019). Despite being considered demographically “recovered”, a previous study comparing the Kirtland’s warblers to their closest relatives showed evidence for recent inbreeding and a high burden of deleterious variation (Calderón et al., 2024). Here, we use whole genome sequencing of contemporary (2000s) and historical samples (early 1900s) to understand the role of recent declines in shaping genetic load. We complement these analyses with forward-in-time simulations informed by the species’ demographic history and genomic structure to validate our results and evaluate how missing data and sample size influences the ability to detect changes in genetic load.

**Figure 1:**
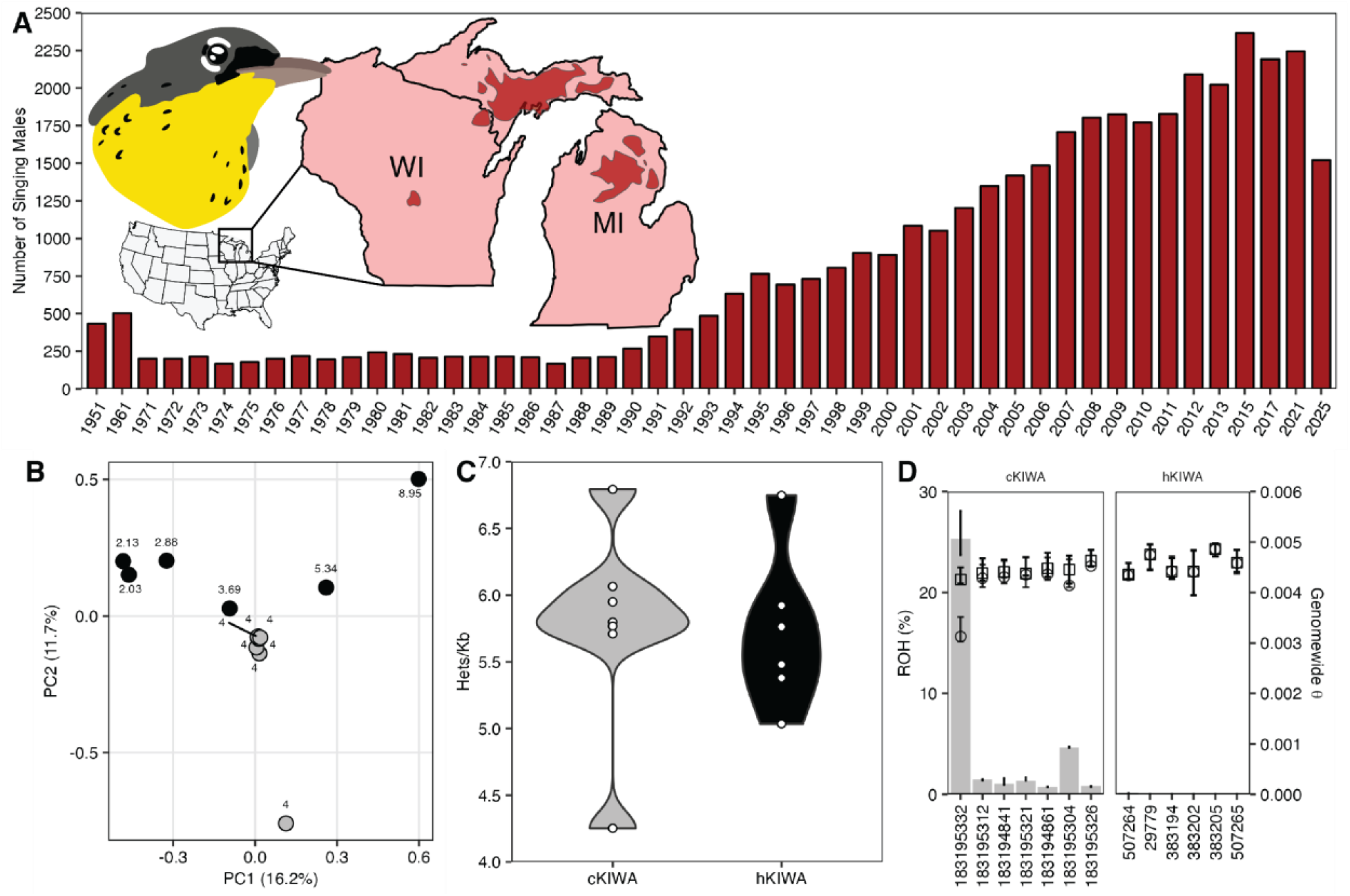
Population trends, population structure, and genetic diversity patterns before (hKIWA) and after (cKIWA) strong bottleneck. **A)** The inset map shows the Kirtland’s warbler breeding distribution (darker shaded polygons) across the states of Michigan and Wisconsin, with the core breeding range vastly concentrated in the lower peninsula of Michigan (Fink et al., 2025). Census counts are reported as the number of singing males and were conducted across Michigan every decade from 1951-1971, annually from 1972-2015, and then every four years from 2017-2025. **B)** PCA shows little to no population structure within each of the historical (black) or contemporary (gray) populations. The numbers beside each point indicate the sequencing coverage. **C)** Local heterozygosity estimates across samples are reported as the weighted mean heterozygous sites per 1000 bp weighted by the number of validated sites. **D)** Shows the percentage of genome within ROH. The secondary y-axis reports Watterson’s θ when including ROH (circles) or excluding ROH (squares). ROHan reports confidence intervals as the minimum and maximum values from converging MCMC chains.

## METHODS

### Sampling and Sequencing

To compare genetic diversity metrics pre- and post-bottleneck, our samples included both historical and contemporary *S. kirtlandii* individuals. Tissues from the historical samples were collected from the toe pads of 10 museum specimens (all male) at the American Museum of Natural History (AMNH). Historical individuals were originally caught and collected in their wintering grounds in the Bahamas or during the breeding season in Michigan between 1884-1922. The contemporary samples included 7 males originally sampled on their breeding grounds in Michigan in 2006. These samples were previously included in (Baiz et al., 2021) and (Calderón et al., 2024) (**Table S1**).

Historical sample digestion and DNA extraction were performed in dedicated historical DNA facilities at the AMNH following contamination-controlled procedures (Fulton, 2012; Knapp et al., 2012). Toe-pad subsamples were minced and briefly washed in 70% ethanol followed by two washes in ultrapure water, each for 2–3 h, to reduce surface contaminants and residual preservatives. DNA was extracted using a modified Qiagen− DNeasy Blood & Tissue protocol optimized for degraded DNA, with overnight digestion in 180 µL buffer ATL and 30 µL proteinase K (20 mg/mL) at 56°C on a thermomixer at 850 rpm. Lysates were then combined with 200 µL buffer AL and 220 µL ethanol and purified following the protocols for the Qiagen− DNeasy Blood & Tissue kit, except that Qiagen− QIAquick spin columns were used in place of standard DNeasy columns to improve retention of the short DNA fragments characteristic of historical material. DNA was recovered using a double elution in 100 µL TE buffer to maximize yield. Library preparations were performed at Penn State in a standard laboratory using TruSeq Nano DNA Library Preparation kit, with one important modification. Given the degraded nature of historical DNA, we omitted the size selection step and instead used undiluted SPB (sample purification beads) to capture DNA fragments of any size (Wood et al., 2023). We did not recover sufficient DNA from sample #383195, so this individual was dropped from any further analysis.

The remaining libraries were sequenced at Penn State’s Genomics Core Facility. A Library tapestation showed varying amounts of adapter dimer on all samples resulting from the library preparation and low sample input. The samples with the highest adapter dimer, #1571 and #26747, were omitted and the remainder of the 7 samples were pooled. These were then size selected using a BluePippin to discard excess adapter dimer. We sequenced the pooled samples on a single lane of Illumina NextSeq 2000 150bp paired end with P2 chemistry.

### DNA Damage, Variant Calling, and Genotype Likelihoods

We bioinformatically processed 7 historical and 7 contemporary *S. kirtlandii* samples. We also included 12 previously sequenced individuals from closely related sister species, *Setophaga citrina* (n=5) and *Setophaga ruticilla* (n=7), for genotype polarization for our downstream analysis. We used AdapterRemoval V2 (Schubert et al., 2016) to trim adaptors and aligned all samples to the Yellow-rumped Warbler (Myrtle subspecies: *Setophaga coronata coronata*) chromosome-level reference genome (PRJNA325157, (Baiz et al., 2021)) using bowtie2 (Langmead & Salzberg, 2012). All sam files were converted to bams, sorted, and indexed with SAMTOOLS V1.18 (Danecek et al., 2021). Duplicates were marked with PicardTools V2.20.9 (https://broadinstitute.github.io/picard). Additionally, to reduce coverage bias, we downsampled contemporary bam files with SAMTOOLS (samtools view -s) by dropping a proportion of reads to achieve an average coverage of 4x across the contemporary *S. kirtlandii* samples. For all samples, coverage was calculated by extracting site depth using samtools depth -a and then taking an average of all sites (excluding the mitochondria, unmapped scaffolds, and the sex chromosome). Sample #749877 was omitted at this stage from further analysis due to low coverage.

To assess post-mortem damage, we ran mapDamage V2.3 (Ginolhac et al., 2011; Jónsson et al., 2013) on the resulting alignment files. We excluded any base quality scores less than or equal to 20 (-Q 20) and considered a read length of 20bp in either read direction (-l 20). For the misicorporation analysis, only the terminal 15 nucleotides at each read end were considered (-m 15). To reduce the effects of post-mortem damage, we rescaled base quality scores at damaged sites using the --rescale flag. This generated .rescaled.bam files that were used in all downstream analysis.

We called variants using GATK4 Haplotype caller (Poplin et al., 2018) and created a .gvcf file for each sample using the -ERC GVCF option. We then combined all samples using CombineGVCFs and genotyped them with GenotypeGVCFs. We subsequently filtered our vcf by first excluding the sex chromosome, mitochondria, and all unmapped scaffolds. We then filtered for biallelic sites (-m2 -M2 -v snps), and filtered sites based on site quality, including all sites with QUAL>=50. We also filtered genotypes based on read depth and genotype quality by setting genotypes with read depth less than 1 or greater than 27 (likely representing repetitive regions with inaccurate genotype calls) to missing as well as those with genotype quality less than 20. We polarized genotypes by defining derived sites as those where all outgroup samples (*S. ruticilla* and *S. citrina*) were homozygous for the reference but at least one *S. kirtlandii* sample contained a heterozygous alternate. After polarization, we retained a total of 8,250,438 derived sites.

Given the low coverage of the historical and downsampled contemporary samples, we also generated genotype likelihoods and minor allele frequencies with ANGSD V0.938 (Korneliussen et al., 2014) using the following presets: -GL 1, -doMajorMinor 1, -doMaf 1 -doCounts 1, - minMapQ 30, -minQ 20, -remove_bads 1, -only_proper_pairs 1, -uniqueOnly 1, -C 50, -baq 1. Similarly, we polarized sites by comparing the frequency of major and minor alleles in the outgroup versus ingroup samples. Specifically, we ran ANGSD separately on the outgroup and ingroup species. The outgroup comprised *S. citrina* and *S. ruticilla*, while the ingroup consisted of both contemporary and historical *S. kirtlandii* samples. We defined derived sites as those where the outgroup was fixed for the reference allele and intersected these with ingroup sites that were segregating for a non-reference minor allele. To assess allele frequency changes, we re-ran ANGSD on the *S. kirtlandii* samples, analyzing contemporary and historical individuals separately. We then filtered sites based on number of individuals, keeping sites where 3 or more individuals contributed to allele frequency estimates. After intersecting these with our list of derived sites, we retained 8,201,989 sites. To account for potential post-mortem damage, we further filtered out sites where the major and minor alleles were transition pairs (T/C, C/T, G/A, or A/G) which resulted in a total of 2,892,290 sites.

### Population Structure and Genetic Diversity

We explored population structure by conducting a PCA analysis with PLINK (Purcell et al., 2007). Using our filtered VCF, we first performed linkage pruning to reduce variable correlations resulting from linkage disequilibrium. We ran PLINK on 50kb windows, a window step of 10bp, and an r^2^ threshold of 0.1. After linkage pruning, we re-ran PLINK to create our PCA using our pruned data. With the exception of the --indep-pairwise flag, we specified the same parameters as before as well as set the --pca flag and included our prune.in file with the --extract option.

Next, we assessed changes in heterozygosity and inbreeding over time by calling ROH using ROHan, a probabilistic method that jointly estimates genetic diversity while accounting for sequencing errors from low coverage data and postmortem damage (Renaud et al., 2019). ROHan estimates local and global heterozygosity rates using a log-likelihood approach weighted by site and genome coverage before applying a hidden markov model to identify ROH and calculate Watterson’s θ. We prepared the bam files according to the developer’s recommendations (https://github.com/grenaud/ROHan) and only applied a fail QC flag to remove reads that did not pass basic quality controls (samtools view -b -F 0 × 200). For the historical samples, we implemented the bam2prof option to quantify aDNA damage with -double, -paired, and -classic parameters. This generated a profile of substitution rates at 3’ and 5’ ends that were included when running ROHan (--deam5p, --deam3p). We set the transition/transversion rate to 2.03 (--tstv) and a heterozygosity rate tolerance of 1.5e-04 (-rohmu). Given the low coverage of our data, we ran ROHan on large windows (1Mb). In comparison to our previous study of contemporary samples (Calderón et al., 2024), we found that ROHan slightly underestimated the fraction of the genome in ROH (i.e. FROH) (**Table S2**). This is likely because the use of the 1MB windows limited ROHan’s ability to detect short ROH in sample coverage <5X. Further, to calculate the heterozygous sites per 1000 bp for each sample (Figure 2C), we averaged the local per base pair heterozygosity rates reported by ROHan, weighted by the number of validated sites and then multiplied this weighted mean by 1000.

**Figure 2:**
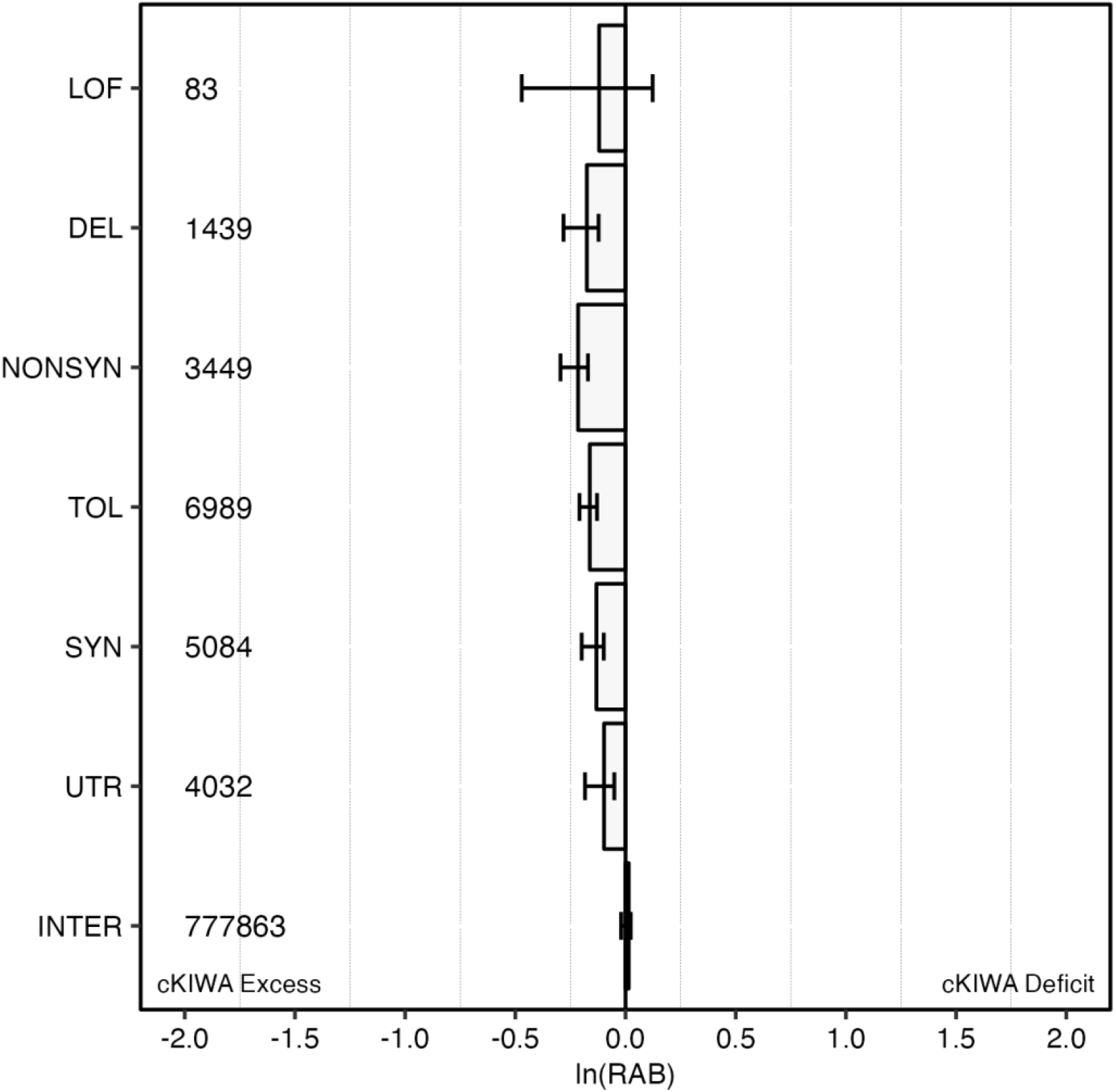
R_A/B_ compares the relative excess of derived alleles between population A (historical *S. kirtlandii*) and population B (contemporary *S. kirtlandii*; cKIWA) across mutational categories and normalized by a set of 10,000 intergenic sites. We report the natural logarithm of R_A/B_ values for a one-to-one comparison, where R_A/B_ < 0 indicates derived alleles in a given set of sites are more frequent in contemporary samples. Whiskers indicate the 2.5% and 97.5% quantiles obtained from running 100 jackknife resamples. The numbers reported on the left indicate the number of sites in each mutational category.

To understand how strong bottlenecks affect genetic diversity, we used Pixy (Korunes & Samuk, 2021) to compare nucleotide diversity between populations. Because Pixy incorporates variant and invariant sites to account for missing data, we generated an all sites vcf with GATK GenotypeVCFs and the -all-sites flag. Using bcftools, we filtered the vcf keeping all biallelic sites (-M2). Additionally, we kept all sites where depth was greater than 10 (INFO/DP>10) and site quality was greater than or equal to 30 (QUAL>=30). To calculate a genome-wide nucleotide diversity estimate, we ran Pixy with a window size of 1 and then manually calculated pi by dividing the reported sum of differences by the sum of total comparisons. To visualize, we re-ran Pixy using a window size of 100kb.

### Deleterious Mutational Load Across Time

We quantified the change in mutational load using functionally annotated variants from a database that was previously constructed (Calderón et al., 2024) with the Myrtle Warbler mywagenomev2.1 assembly (Baiz et al., 2021) and the program SIFT4G (Vaser et al., 2016). Briefly, SIFT4G predicts how amino acid changes will impact protein function by evaluating how conserved the position is among homologous protein sequences. It classified mutations as noncoding (n=281,587), synonymous (n=337,158), nonsynonymous (n=207,513), or loss-of-function (n=5,239). For synonymous and nonsynonymous mutations, SIFT further predicted whether they had a tolerated (n=450,515) or deleterious (n=85,028) effect on protein function. To obtain a list of derived mutations, we overlapped our annotated variants with derived sites in our polarized vcf. For each mutational category, we tracked changes in mutational load by computing R_A/B_ using a custom script. R_A/B_ is a statistic that measures the excess or deficit of derived alleles of a specific category (e.g, loss-of-function) between two populations while normalizing over neutral sites. It is defined as 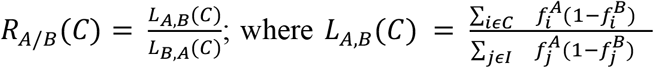 2015). We assigned population A as the historical individuals, population B as the contemporary individuals, and normalized against a set of 10,000 independent intergenic sites. To create quantile intervals, we ran 100 jackknife resamples, randomly subsampling 30% of the data with each jackknife replicate.

### Simulations

To test whether the R_A/B_ estimates reflected a true signal of genetic load shifts, we replicated this analysis with simulations. Using SLiM V5.1 (Haller et al., 2026), we conducted Wright-Fisher simulations by assuming a randomly mating hermaphroditic population. To simulate a demographically realistic population, parameters were set using molecular Ne inferences ((Wilson et al., 2012) using an initial population of 700 and a burn of 10,000 generations. After burn-in, we simulated an instant contraction to 259 individuals for 275 generations followed by a secondary contraction to 161 individuals. This secondary contraction represents the strong bottleneck that occurred between 1940-1950.

For these *in silico* genome simulations, we constructed the genome structure using the Myrtle Warbler autosomal genome, importing all exons and intergenic regions from the available gene annotations and using only transcripts on the positive strand. This resulted in a total of 13,492,811 exonic base pairs. We allowed only neutral mutations to occur in intergenic regions. Within exons, both neutral and deleterious mutations occurred at a 2:3 ratio, respectively. Neutral mutations were sampled from a fixed distribution while deleterious mutations were drawn from a gamma distribution with a mean *s* = -0.0131 and shape parameter α = 0.186 (Kim et al., 2017; Kyriazis et al., 2021). We set mutations to arise at a rate of 4e-9 per site per generation, which was the estimated mutation rate for another songbird, the Collared Flycatchers (Smeds et al., 2016). We set the average recombination rate to 3.1cM/Mb or 3e-8 per site per recombination (Kawakami et al., 2014). We explored the effect of dominance by varying the dominance coefficients (*h*) for deleterious alleles from completely recessive to completely dominant, *h* ∈ {0, 0.25, 0.5, 1.0}. Lastly, we simulated two sampling events to represent our two datasets with the first sampling event occurring 25 generations prior to the bottleneck and the second sampling event at 28 generations after the bottleneck. To understand how sample size affects the ability to detect changes in genetic load, we varied our sample size. First, we ran simulations with increased sample sizes N_pre_=20, N_post_=20 to obtain the expected change in genetic load when not constrained by sample size. Then we re-ran simulations using a lower sample size to represent the true circumstances (N_pre_=6, N_post_=7) in our empirical data.

This resulted in a total of eight simulations replicated 300 times for each dominance scenario to ensure sufficient moderately to strongly deleterious mutations for statistical analysis. Using the resulting VCFs, we further explored the effects of missingness on R_A/B_ estimates by introducing missing genotypes using Empirical Genotype Generalizer for Samples (EGGS) (Smith et al., 2026). When the “-b” option is set, EGGS uses patterns of missingness from an empirical VCF (e.g., per-individual and per-site missingness rates) to parameterize a beta-distribution and randomly introduce similar patterns of missingness in a simulated VCF. We computed R_A/B_ across all simulated VCFs in four mutational categories: neutral, nearly neutral, weakly deleterious, mildly deleterious, moderately deleterious, and strongly deleterious. Neutral mutations were defined as exonic variants with selection coefficient of *s* = 0. Nearly neutral mutations were defined as those with -0.0001 < s < 0, weakly deleterious as -0.001 < s ≤ -0.0001, mildly deleterious mutations as -0.01 < s ≤ -0.001, moderately deleterious as -0.1 < s ≤ -0.01, and strongly deleterious mutations as those with s < -0.1.

## RESULTS

### Sequencing and Variant Calling

Sequencing of historical *S. kirtlandii* samples resulted in a range of low coverages between 2-9X. Contemporary samples were previously sequenced to an average coverage of 18X but downsampled in this study to 4X to reduce potential batch effects. All other *S. ruticilla* and *S. citrina* samples used for genotype polarization were previously sequenced to 17X (**Table S1**).

To assess postmortem degradation, we ran MapDamage on both historical and contemporary samples. In both contemporary and historical samples, the frequency of misincorporated bases was at 0 for the first 3-4 positions near fragment ends, likely resulting from adapter removal (**Figure S1A, B**). In historical samples, the frequency of misincorporated bases declined at positions furthest from the terminal ends, a pattern expected from DNA damage. Additionally, there was a slightly higher frequency of G→A substitutions on the 3’ ends in the historical samples (**Figure S1A**). This pronounced effect on the 3’ ends could be due to library preparation biases where the tagmentation process may have a higher efficiency for repair on the 5’ ends, sequencing platforms reading strands in one direction, or slight differences in DNA fragmentation. In the contemporary samples, the frequency of misincorporated bases was consistent regardless of position from fragment end and this was expected given that these samples were not affected by postmortem degradation.

### Population Structure and Genetic Diversity

Our principal components analysis showed little to no population structure in each of the contemporary and historical samples (**Figure 1B**). Principal component 1 explained 16.2% of the variation and separated samples based on sequence coverage. Historical samples separated most clearly from low to higher coverage across PC1 while contemporary samples all with an average coverage of 4X clustered at the center of PC1. Samples were further separated across PC2 which explained 11.7% of variation. Except for one, all contemporary samples clustered at the center of PC2, while the historical samples showed slightly higher variability along this axis. This pattern likely reflects the spectrum of genetic variation across samples, which is shown more clearly in **Figure 1C**. Overall, genome-wide heterozygosity patterns across samples was similar between populations with an estimated 5.76 hets/kb in the contemporary population and 5.72 hets/kb in the historical (**Figure 1C**).

Next, we characterized the patterns of inbreeding using ROHan, a probabilistic method that estimates heterozygosity while accounting for postmortem damage (Renaud et al., 2019). We ran ROHan using 1Mb windows to increase ROH detection. ROHan detected little to no inbreeding in historical genomes whereas the contemporary samples showed varying FROH ranging from 0.74-25% (**Figure 1D, Table S2**). FROH estimates in contemporary samples were similar to those reported in our previous study which used a different analytical tool, GARLIC (Szpiech et al., 2017)(**Table S2**). Despite differences in FROH and heterozygosity between populations, genome-wide Watterson’s θ outside of ROH was similar across all samples. This was consistent with genomewide nucleotide diversity estimates as calculated by pixy (**Figure S2**) (Korunes & Samuk, 2021). When including ROH, however, Watterson’s θ noticeably decreased when FROH was substantially high (**Figure 1A**).

### Genetic Load Across Time

#### Empirical Results

To understand how population bottlenecks affect genetic load in historically small populations, we calculated R_A/B_, a statistic that compares the relative excess of derived alleles between two populations while normalizing against a set of putatively neutral sites (Xue et al., 2015). Our estimates indicated an overall excess of derived alleles in the contemporary samples across all mutational categories (**Figure 2**). Given the uniformity in our results, we suspected that calling variants may have potentially introduced a bias in our data. To investigate this possibility, we also calculated R_A/B_ from minor allele frequencies generated from genotype-likelihoods in ANGSD. Further, to determine if this potential bias originated from our putatively neutral sites, we normalized over both intergenic and four-fold synonymous sites. Results from minor allele frequencies revealed a similar pattern that was originally identified with called genotypes, and this was consistent regardless of whether we used intergenic or four-fold synonymous sites for standardization (**Figure S3A, B**). Furthermore, this result persisted when removing transition pairs potentially resulting from DNA damage (**Figure S3C, D**), suggesting the finding was robust.

We used forward-in-time simulations to explore the expected changes in genetic load for a population of similar demographic history as the Kirtland’s warblers. We ran simulations using SLiM5 (Haller et al., 2026) under a Wright-Fisher model for four dominance scenarios: complete recessiveness, partially recessive, additive, and completely dominant. We simulated the long-term and recent population history of the Kirtland’s Warblers and constructed an identical genomic structure using annotations from the reference genome (Baiz et al., 2021). Across all dominance scenarios, our simulations indicate that the relative excess of mutations depended on both selection and their dominance (**Figure 3A**). Specifically, recessive mutations did not show any notable deviations from 0. As dominance increased, however, mildly to strongly deleterious mutations (s≤-0.001) accumulated in the post-bottleneck populations. Although accumulation occurred in the strongly deleterious category (s<-0.1), we interpret this category cautiously given that the average number of mutations per replicate was only 3 (**Figure 3C**).

**Figure 3:**
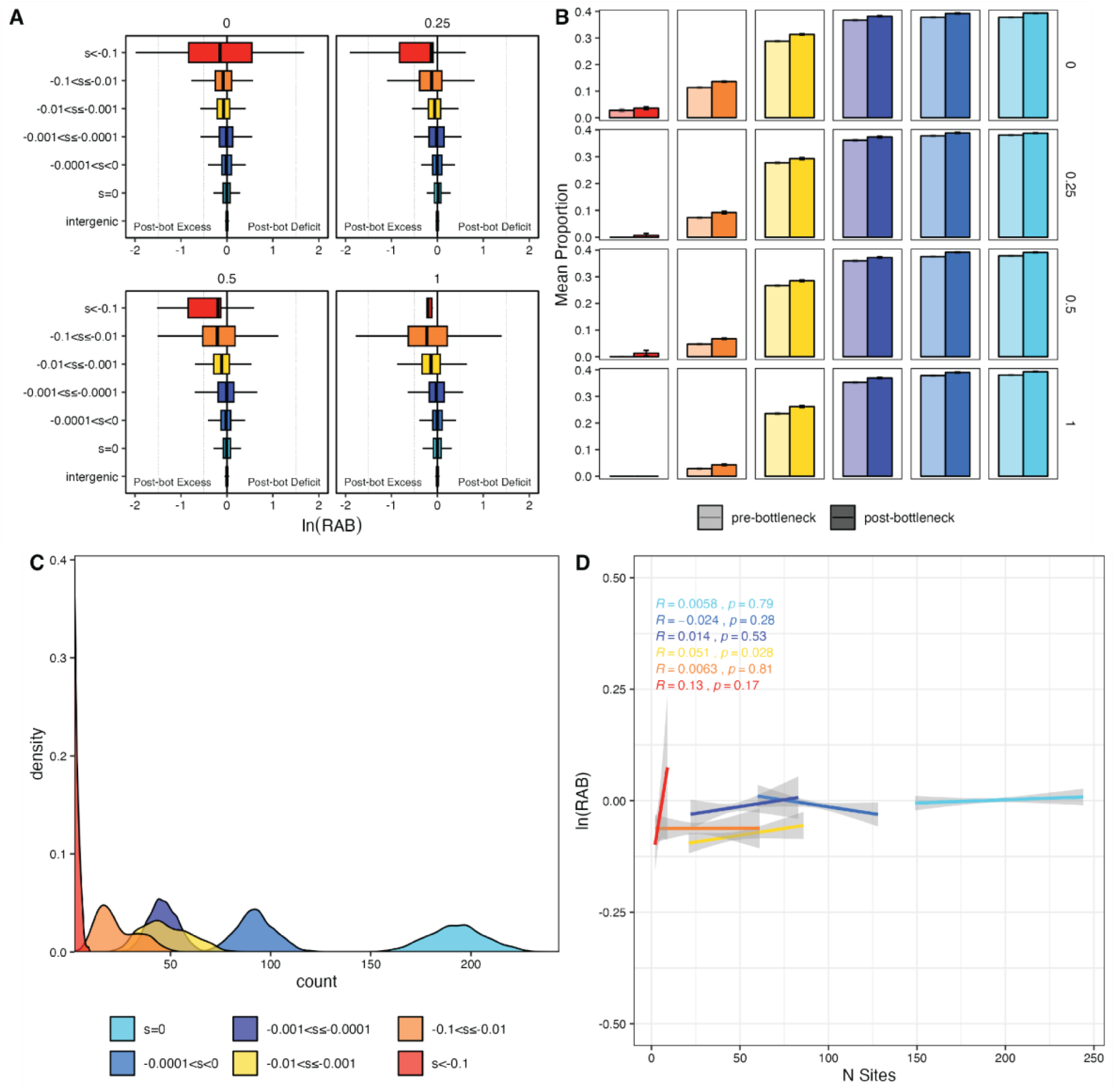
Changes in genetic load and composition across dominance coefficients (N_pre_=20, N_post_=20) **A)** box plots show the distribution of R_A/B_ estimates across all replicates where the median is denoted with a black vertical line in the boxplot. **B)** shows the mean proportion of derived homozygotes for each mutational category and its respective dominance in pre- and post-bottleneck populations. The whiskers represent standard error and each mutational category was plotted on an unscaled y-axis for better comparison. **C)** distribution of mutations across all 300 replicates. Dominance coefficients here have been collapsed but general distribution patterns are retained under each dominance scenario. **D)** Correlation between R_A/B_ estimates and number of mutations across all dominance scenarios and replicates. Labels indicate Pearson’s correlation coefficients and their associated p-values. Colors indicate mutation categories and are consistent across all four panels.

To understand how bottlenecks alter the composition of genetic load, we tracked the number derived genotypes, specifically focusing on homozygotes (**Figure 3B**). For each mutation class, dominance coefficient, and replicate, we calculated the proportion of derived homozygotes in each population. Specifically, the proportion of derived homozygotes was computed by dividing the number of derived homozygotes by the total number of derived genotypes (i.e., the sum of derived homozygotes and derived heterozygotes), then summarized by taking the average across replicates. Although we did not observe any clear differences in neutral to weakly deleterious categories (s > -0.001), the mean proportion of derived homozygotes in mild (-0.01 < s ≤ -0.001) and moderately deleterious mutations (-0.1 < s ≤ -0.01) increased in post-bottleneck populations across all dominances. Interestingly, no change in derived homozygotes was observed for strongly deleterious (s<-0.1) recessive mutations. A slight increase was observed in partially recessive and additive mutations. That said, we again exercise caution in interpreting this category due to the small number of mutations produced in each replicate (**Figure 3C**).

Given the differences in the distribution of the number of mutational sites across simulations (**Figure 3C**), we asked if the number of mutations of a specific category could drive the estimates observed in our simulations. Without taking dominance into account, Pearson’s correlation coefficients indicated mostly weak but positive correlations between R_A/B_ and the number of sites of a specific mutational category. However, only the mildly deleterious category resulted in a significant association (p-value > .05) (**Figure 3D**).

Lastly, we explored how missing data and sample size could affect our capacity to detect changes in genetic load. First, we ran simulations using a high sample size (N_pre_=20, N_post_=20) and then introduced patterns of missing genotypes that we would expect to find in empirical data. Our results show that missing data disproportionately affected estimates in the moderately to strongly deleterious categories (**Figure 4A**). In the moderate category, missingness inflated the accumulation of mutations in post-bottleneck populations. In the strongly deleterious category, it vastly inhibited our ability to detect any changes in genetic load. We then ran simulations with a lower sample size identical to our real sampling scheme (N_pre_=6, N_post_=7). Like before, our results showed that reduced sample sizes disproportionately affected estimates in the moderate and strongly deleterious categories (**Figure 4B**). Namely, reduced sample sizes either inflated accumulation patterns or inhibited our ability to detect accumulation. Taken together, our results show that sample size, missingness, and the number of mutations can obscure signals of changes in genetic load.

**Figure 4:**
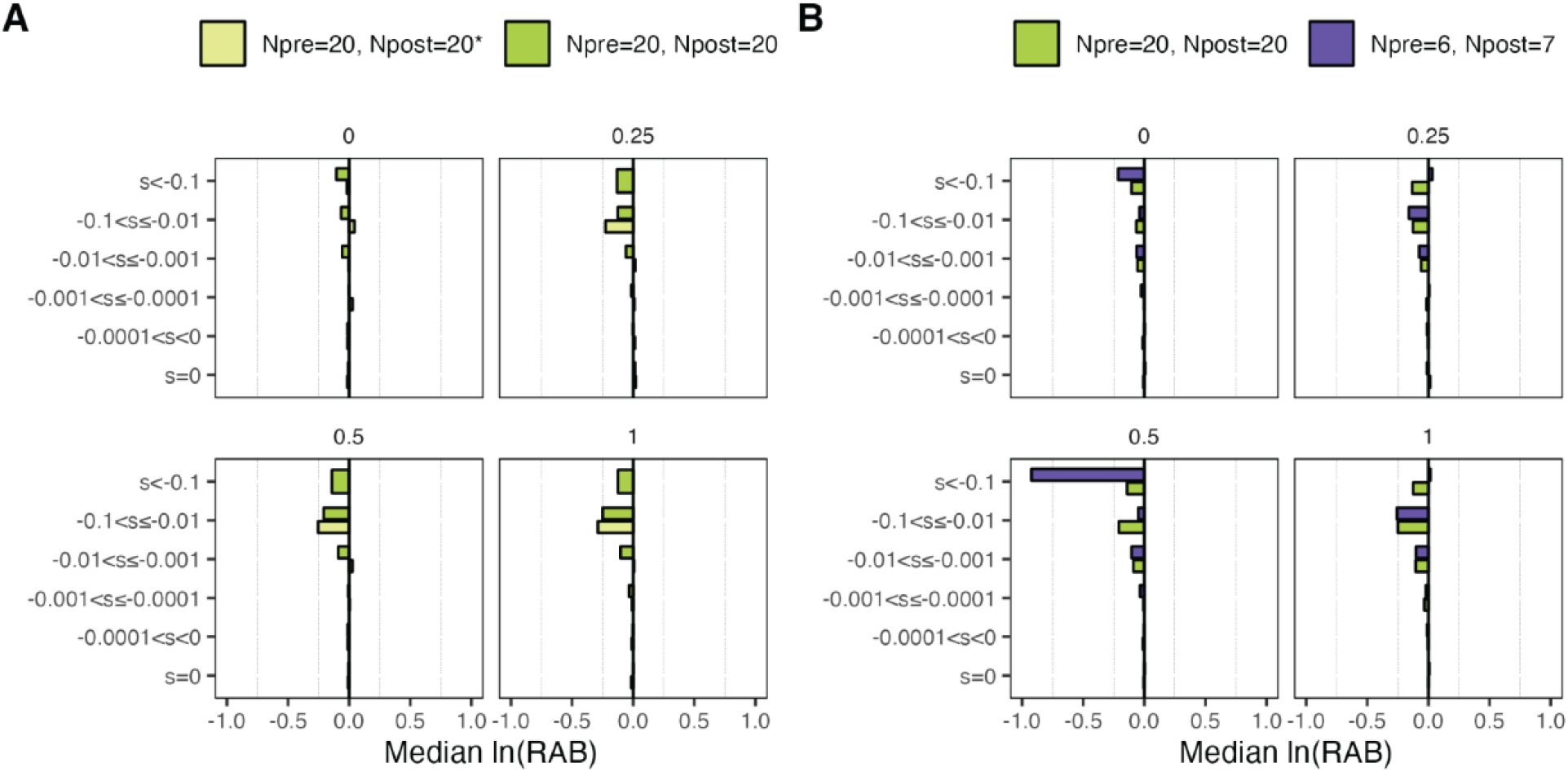
Effects of missingness and sample size on capacity to detect changes in genetic load. **A)** shows the median R_A/B_ estimates across replicates when missing data is introduced (light green) and without any missing data (dark green). **B)** compares estimates in simulations with higher sample sizes (green) versus low sample sizes (purple).

## DISCUSSION

In this study, we show that despite an ancestrally small population, Kirtland’s warbler populations were large enough to avoid close inbreeding prior to the 1940s and that changes in genetic load were likely driven by more recent demographic declines. Contemporary samples showed an accumulation of genetic load across all mutational categories, a pattern that also appeared in our simulations. Overall, our simulations indicate that bottlenecks can indeed lead to accumulation of deleterious genetic load in long-term small populations and in a relatively short time frame (∼28 generations). Previous studies have found similar patterns of accumulation of mildly deleterious mutations (Dussex et al., 2021; Grossen et al., 2020; Xue et al., 2015). But contrary to these studies, which also document the purging of strongly deleterious mutations, both our empirical data and simulations showed the opposite effect in loss-of-function mutations and mutations with s < -0.1 suggesting a reduced purifying response during this recent population decline. While this result is surprising given the frequent documentation of purging in small populations (van der Valk et al., 2019), a similar pattern has also been observed in Chatham Island Black Robins despite experiencing a prolonged and severe bottleneck (Kennedy et al., 2014).

Unlike the Black robins, which showed reduced juvenile survival with increased inbreeding coefficients, our samples showed no clear phenotypic signs of overt inbreeding depression. Such observations, however, are challenging for a neotropical migratory songbird, though opportunistic empirical field observations indicate the potential for reduced hatching rates. Moreover, recent field surveys documented at least one case of albinism in Kirtland’s Warblers nestlings. Albinism is a rare mendelian autosomal recessive condition that leads to disruptions in the melanin pathway and has been linked to consanguinity in humans (Bakker et al., 2022; Farooq et al., 2025; Gamella et al., 2013). Albinism is also reported in non-human species and may be more common among domesticated, captive bred, and small isolated populations. In great reed warblers (*Acrocephalus arundinaceus*) for example, the occurrence of partial albinism was highest following population colonization in Sweden, reported at 4.5% (Bensch et al., 2000). While only a single observation out of 39 surveyed nests, the frequency of albinism and other inbreeding-related traits in the Kirtland’s warblers warrants further investigation. Nevertheless, these observations imply that inbreeding depression could operate cryptically and pose a complication for this population’s ongoing management. More generally, it also suggests that conservation geneticists should be cautious about overemphasizing the concept of purging in other historically small populations

While our results are consistent with the limitations to purging, some caveats remain— highlighting how sample size, missing data, and number of mutations may also influence the ability to detect changes in genetic load. For example, we show that missing data and low sample size most commonly inflates signals of genetic load accumulation in the most deleterious category. Further, we also showed that the number of sites could affect R_A/B_ estimates, so we recommend exercising caution when interpreting such statistics. In situations where sample size is lacking, the use of genetically informed simulations can supplement empirical inference by establishing null expectations under similar demographic scenarios. Forward in time simulations are increasingly being used to explore conservation-relevant questions but we recognize that they may be limited to some key assumptions. First, the distribution of fitness effects (DFE) describes the selective effects of new mutations and their relative frequencies. The DFE is an essential component for understanding the quantitative genetic variation underlying complex traits and the consequences of reduced population size. But there remains considerable debate about DFE estimates even in well-studied populations such as humans (Boyko et al., 2008; Eyre-Walker et al., 2006; Kardos et al., 2021; Kim et al., 2017; Pérez-Pereira et al., 2022). Additionally, as far as we are aware, only one study has attempted to estimate the DFE outside of primates in a non-model organism, the vaquita porpoise (Robinson et al., 2022). Other estimates from model organisms like yeast, *Drosophila*, and mice suggest that DFEs vary substantially between species (Huber et al., 2017; Kyriazis et al., 2023).

Second, the mutational target size also depends on the number of genes or exons, their lengths, the mutation rate, and the proportion of deleterious to neutral mutations arising in the population. The genome structure used for Kirtland’s Warblers was derived from a reference annotation; however, only exons annotated on the positive strand were included in the simulations. As a result, the total number of coding sites represented in the model is reduced relative to the full genome, which may decrease the number of mutations that arise in coding regions in our simulations. Estimates suggest that mutation rates only vary moderately between bird species. For this study, we used mutation rates estimated in collared flycatchers (4 × 10^−9^ per site per generation; (Smeds et al., 2016)) which is comparable to estimates in zebra finch (5 × 10^−9^ per site per generation; (Prentout et al., 2025)). Overall, our deleterious mutations appeared at a rate of U=0.064. For comparison, U_*Drosophila*_ = 1.2 (Haag-Liautard et al., 2007), so our estimate is 18.5 times lower than that of *Drosophila* (U_*Drosophila*_/U_*Setophaga*_). This conservative proportion of deleterious mutations may underestimate the effects of genetic load in the real world.

Recombination can also influence the purging of deleterious mutations. In our simulations we used an average genome-wide recombination of rate 3.1cM/Mb (Kawakami et al., 2014). In reality, low recombinant regions create stronger linkage, thereby intensifying Hill-Robertson effects and is further exacerbated by inbreeding as the effective recombination rate decreases (Comeron et al., 2008). Lastly, studies on the percentage of lethal mutations are needed in non-model organisms to understand the relative contribution of large-effect alleles to fitness. While we did not simulate any autosomal recessive lethal mutations, they are likely to occur as they do in other species such as *Drosophila* (∼1.6 per individual), humans (1-2 per individual;(Gao et al., 2015), and other wild bird populations like the ruff (Philomachus pugnax;(Küpper et al., 2016) and the Scottish population of Red-billed Choughs (*Pyrrhocorax pyrrhocorax*;(Trask et al., 2016).

In conclusion, the apparent lack of purging in Kirtland’s Warblers may reflect a combination of long-term small Ne as well as methodological biases. Although purging of strongly deleterious alleles is theoretically expected under sustained inbreeding, our study adds to evidence that its efficacy can be limited under certain demographic and selective regimes. Overall, our results are consistent with population genetic theory predicting that in small populations, where drift can overwhelm selection, mild and moderately deleterious mutations can rise to appreciable frequencies. Like other bottlenecked populations, Kirtland’s warblers had an excess of mildly to moderately deleterious mutations. This excess likely resulted from the accumulation of drift load, which increases as populations decline. Historically small populations are particularly vulnerable to increasing drift load given that they already possess low genetic diversity to begin with. This is particularly relevant to this species since our previous study showed that they have been historically small since their population diverged from their closest relatives ∼1 million years ago (Calderón et al., 2024). Small populations are also expected to have lower mean fitness due to the expression of partially recessive deleterious alleles (Kardos et al., 2021). Indeed, our simulations indicated the greatest accumulation occurred in partially recessive to dominant mutations. In populations amenable to genetic management, drift load can be reduced by increasing population connectivity or through sustained genetic rescue given a similar genetic load from the donor population (Aguilar-Gómez et al., 2025). Due to the lack of population structure in this species, genetic management is unfeasible. Consequently, to protect this population from the possibility of mutational meltdown, conservation efforts should prioritize maintaining stable population sizes through habitat management both in the wintering and breeding grounds.

## ACKNOWLEDGMENTS

We would like to acknowledge the American Museum of Natural History for contributing historical Kirtland’s warbler samples to this project. Permits for work were supported by the USGS (Master banding permit #24043), the Pennsylvania Game Commission (#46121), and the Pennsylvania Department of Conservation and Natural Resources. We thank Dr. Leilton Luna for his guidance on library preparations, Dr. Matthew Williams for his guidance on bioinformatic pipelines, and Thomas Q. Smith for his help with simulating missing data. Computations were performed using PSU’s Institute for Computational Data Sciences Roar Supercomputer. This work was supported by the National Institute of General Medical Sciences of the National Institutes of Health award number R35GM146926 (ZAS) and the Science Achievement Graduate Fellowship (A.M.C.). This material is based upon work supported by the National Science Foundation Graduate Research Fellowship Program under Grant No. DGE1255832 to A.M.C. and DEB Grant No. 2131469 to D.P.L.T. Any opinions, findings, conclusions, or recommendations expressed in this material are those of the author(s) and do not necessarily reflect the views of the National Science Foundation.

