## Supplementary Information for "Historically Small Population Size Limits Purging of Deleterious Mutations in a Conservation-Reliant Species, the Kirtland’s Warbler"

**Table S1**: Sample information and average autosomal coverage.

| Collection | Species | Sample | Seq | Sex | Prep | Loc | Date | Cov | Accession |
|---|---|---|---|---|---|---|---|---|---|
| CUMV | ruticilla | 163 | Seq[b] | M | Tissue | NY, US | 2003-06-26 | 15.49 | SAMN16870856 |
| CUMV | ruticilla | 1049 | Seq[b] | M | Tissue | NY, US | 2003-07-07 | 17.69 | SAMN16870855 |
| CUMV | ruticilla | 1940 | Seq[b] | M | Tissue | NY, US | 2007-05-15 | 18.28 | SAMN16870857 |
| CUMV | ruticilla | 4056 | Seq[b] | M | Tissue | NY, US | 2013-05-17 | 17.2 | SAMN16870858 |
| D.Toews | ruticilla | 284029323 | Seq[b] | M | Blood | NY, US | 2017-06-09 | 16.53 | SAMN16870853 |
| D.Toews | ruticilla | 283030179 | Seq[c] | M | Blood | NY, US | 2020-06-19 | 18.17 | SAMN40160019 |
| D.Toews | ruticilla | 283030100 | Seq[c] | M | Blood | PA, US | 2020-06-03 | 19.61 | SAMN40160020 |
| CUMV | citrina | 262 | Seq[b] | M | Tissue | NY, US | 2003-08-24 | 18.09 | SAMN16913428 |
| CUMV | citrina | 2871 | Seq[b] | M | Tissue | NY, US | 2009-08-25 | 12.56 | SAMN16913429 |
| D.Toews | citrina | 283030065 | Seq[c] | M | Blood | PA, US | 2020-05-22 | 18.67 | SAMN40160016 |
| D.Toews | citrina | 283030083 | Seq[c] | M | Blood | PA, US | 2020-05-30 | 17.87 | SAMN40160017 |
| D.Toews | citrina | 284029450 | Seq[c] | M | Blood | PA, US | 2019-05-25 | 18.27 | SAMN40160018 |
| UWBM | citrina | 104049 | Seq[b] | M | tissue | GT | 2002-01-14 | 3.94 | SAMN16953418 |
| UWBM | citrina | 107057 | Seq[b] | M | tissue | NC, US | 2005-06-07 | 9.64 | SAMN16953422 |
| Smithsonian | kirtlandii | 183195332 | Seq[b] | M | Blood | MI, US | 2006-06-10 | 15.2 | SAMN16913439 |
| Smithsonian | kirtlandii | 183194861 | Seq[b] | M | Blood | MI, US | 2006-06-07 | 23.05 | SAMN16913440 |
| Smithsonian | kirtlandii | 183195321 | Seq[b] | M | Blood | MI, US | 2006-06-07 | 24.01 | SAMN16913441 |
| Smithsonian | kirtlandii | 183195304 | Seq[b] | M | Blood | MI, US | 2006-05-24 | 14.02 | SAMN16913442 |
| Smithsonian | kirtlandii | 183194841 | Seq[b] | M | Blood | MI, US | 2006-05-27 | 15.06 | SAMN16913443 |
| Smithsonian | kirtlandii | 183195326 | Seq[c] | M | Blood | MI, US | 2006-06-08 | 16.53 | SAMN40160021 |
| Smithsonian | kirtlandii | 183195312 | Seq[c] | M | Blood | MI, US | 2006-05-27 | 20.5 | SAMN40160022 |
| AMNH | kirtlandii | 29779 | Seq | M | Tissue | Bahamas | 1884-02-19 | 2.13 | |
| AMNH | kirtlandii | 383194 | Seq[a] | M | Tissue | MI, USA | 1914-07-06 | 5.43 | |
| AMNH | kirtlandii | 383202 | Seq[a] | M | Tissue | MI, USA | 1915-08-30 | 2.03 | |
| AMNH | kirtlandii | 383205 | Seq[a] | M | Tissue | MI, USA | 1905-06-18 | 2.88 | |
| AMNH | kirtlandii | 507264 | Seq[a] | M | Tissue | Bahamas | 1902-03-20 | 8.95 | |
| AMNH | kirtlandii | 507265 | Seq[a] | M | Tissue | MI, USA | 1904-06-23 | 3.69 | |
| AMNH | kirtlandii | 759877 | Seq[a] | M | Tissue | MI, USA | 1922-06-08 | 1.61 | |

[a]Samples that were newly sequenced in this study.
[b]Samples originally sequenced in Baiz et al. 2021 and re-sequenced in Calderon et al. 2024.
[c]Samples that were originally sequenced in Calderon et al. 2024

**Table S2**: Comparison of inbreeding estimates using GARLIC (Szpiech et al. 2017) and ROHan (Renaud et al. 2019). Estimates from GARLIC were pulled from a previous study (Calderón et al. 2024) in which contemporary samples had a higher coverage. In this study, contemporary samples were down sampled to an average of 4X to reduce sequencing batch effects.

| **Sample** | **Population** | **GARLIC** | **ROHan** |
|---|---|---|---|
| 183195332 | Contemporary | 31.51646254 | 25.2947 |
| 183194861 | Contemporary | 1.89804063 | 0.737619 |
| 183195321 | Contemporary | 2.460903507 | 1.36555 |
| 183195304 | Contemporary | 5.736246705 | 4.63158 |
| 183194841 | Contemporary | 2.320693545 | 1.05263 |
| 183195326 | Contemporary | 1.900475768 | 0.838574 |
| 183195312 | Contemporary | 2.358551116 | 1.46905 |
| 29779 | Historical | | 0 |
| 383194 | Historical | | 0 |
| 383202 | Historical | | 0 |
| 383205 | Historical | | 0 |
| 507264 | Historical | | 0.104822 |
| 507265 | Historical | | 0 |

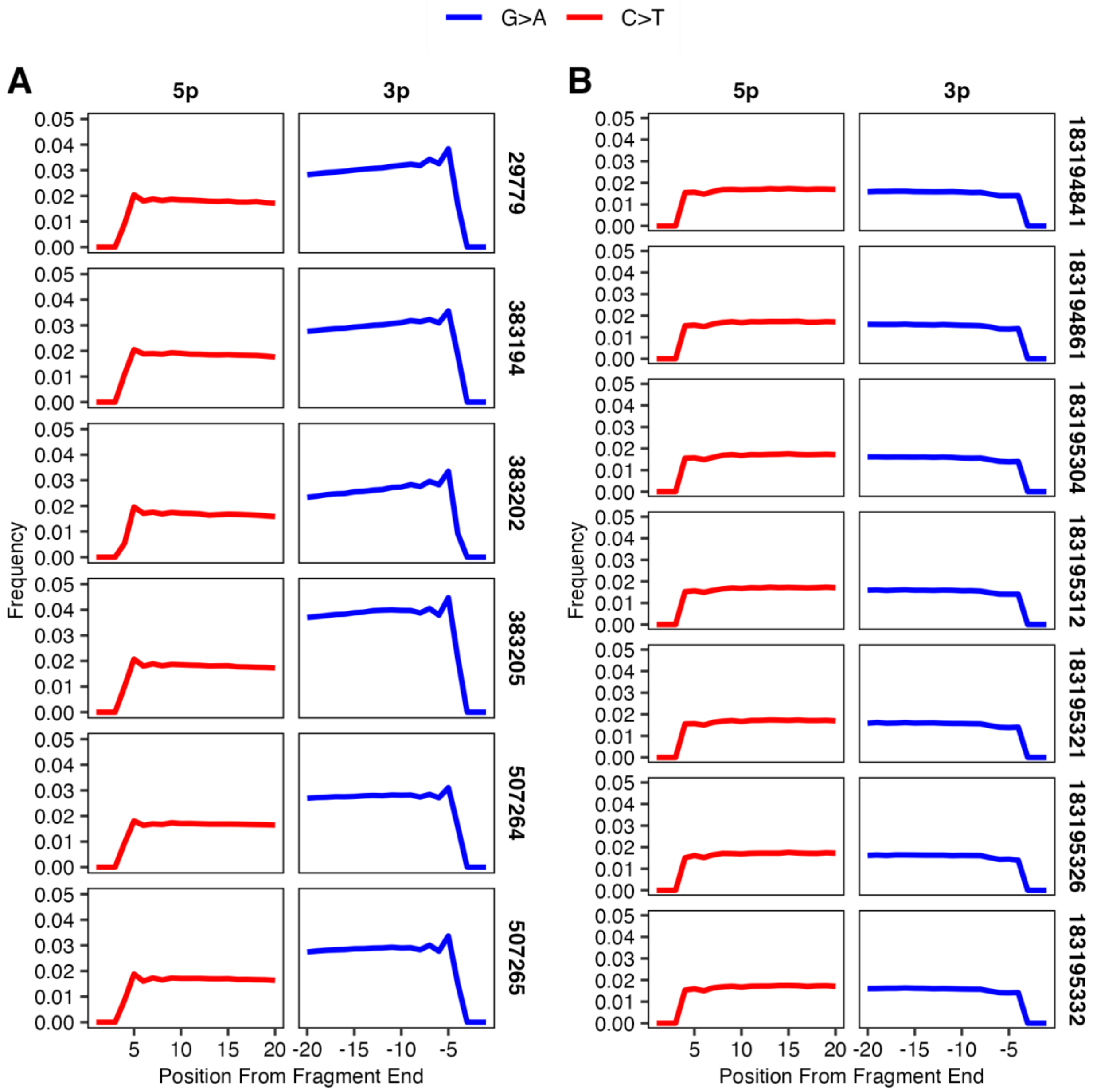


**Figure S1**: Postmortem deamination profiles for **A)** historical samples and **B)** contemporary samples showing substitution rates at varying distances from fragment ends. At the 5′ end on the positive strand, C→T substitution rates are indicated in red. At the 3′ end on the negative strand, G→A substitutions are indicated in blue.

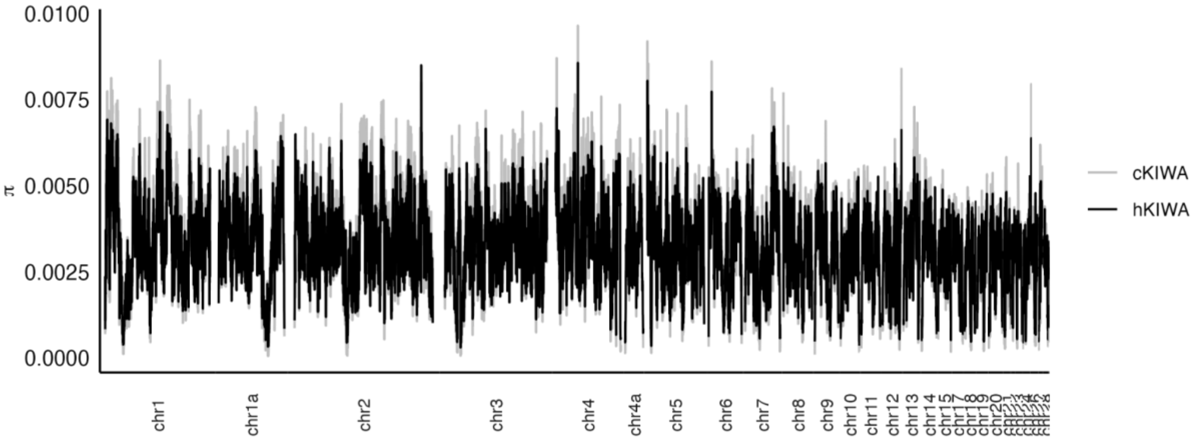


**Figure S2**: Genome-wide π estimates calculated with Pixy in 100kb windows.

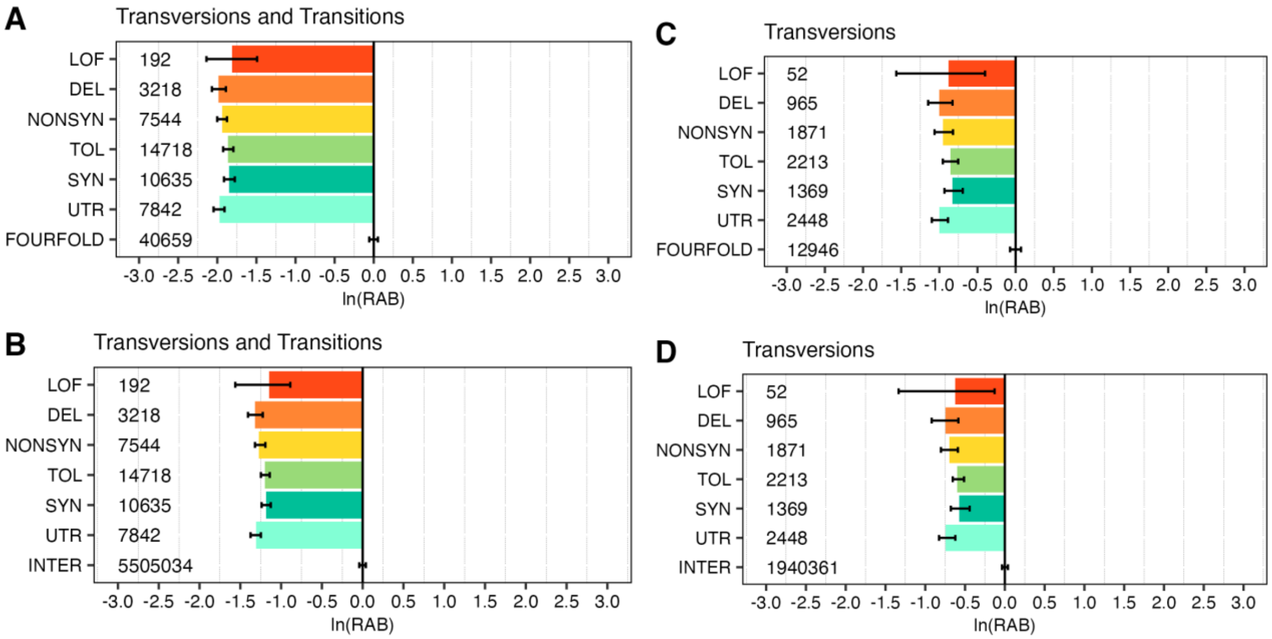


**Figure S3**: $R_{A/B}$ estimates using genotype likelihoods generated with ANGSD. We report the natural logarithm of RAB values for a one-to-one comparison, where $R_{A/B} < 0$ indicates derived alleles in a given set of sites are more frequent in contemporary samples relative to historical samples. Whiskers indicate the 2.5% and 97.5% quantiles obtained from running 100 jackknife resamples. In **A** and **B**, both transversions and transitions are used and normalized by two different sets of 10,000 putatively neutral sites. In **C** and **D**, transitions were filtered out to account for any potentially deaminated bases and again normalized by two different sets of putatively neutral sites.
